# Reactive Oxygen Species Signaling Differentially Controls Wound Healing and Regeneration

**DOI:** 10.1101/2022.04.05.487111

**Authors:** Alanna V. Van Huizen, Samantha J. Hack, Jacqueline M. Greene, Luke J. Kinsey, Wendy S. Beane

**Affiliations:** Department of Biological Sciences, Western Michigan University, Kalamazoo, MI 49008, USA; Department of Hematology, St. Jude Children’s Research Hospital, Memphis, TN, 38105, USA

## Abstract

Reactive oxygen species (ROS), such as hydrogen peroxide, are conserved and critical components of both wound healing and regeneration. Even though millions are affected each year by poor wound healing and an inability to restore functional tissue, how the same ROS-mediated signaling regulates these two different processes is not fully understood. Here, we investigate the role(s) of ROS during planarian wound healing and regeneration. We show ROS accumulate after injury and are required for wound closure (by promoting cytoskeletal movements) and regrowth (by promoting blastema formation). We found that different threshold levels of ROS regulate separate downstream targets to control wound healing (*jun-1*) versus regeneration (*hsp70*). By only manipulating ROS levels, we were able to control which injury-induced program was initiated: failure to close (chronic wound), healing only (no blastema), or full regeneration. Our results demonstrate that healing versus regenerative outcomes are based on differential ROS-mediated gene expression soon after injury. This study highlights ROS signaling as a potential therapeutic means to control wound repair mechanisms in multiple contexts. Therefore, investigating the mechanisms by which ROS control different tissue repair processes will be necessary not only for regenerative medicine but to improve clinical outcomes for chronic wounds and fibrosis.

## Introduction

All organisms have some ability to respond to an injury and repair tissues. Efficient wound response and repair are critical for preventing infection, internal tissue loss, and ultimately, survival. However, tissue repair mechanisms are extremely variable across species and between tissue types. Some organisms can easily regenerate damaged or missing structures after injury, while others replace missing structures with acellular scar tissue that results in a loss of functionality and sometimes disease^1^. Regeneration itself is a highly conserved process; almost every phylum possesses a species capable of some level of functional regrowth^2^. Despite this, not all wounds that heal result in regeneration. Compared to many species, humans are notoriously limited in non-homeostatic tissue renewal. In fact, it has been estimated that in one year alone, for just Medicare patients in the United States, there were over 8 million people affected by chronic wounds with more than $30 billion in care costs across all wound types^3^. Thus, impaired wound healing, chronic wounds and fibrosis-related diseases represent a major health burden^3^.

It is therefore surprising that, despite the many impressive advances made in this field, so much about how different organisms detect and respond to injury to promote functional tissue repair remains unclear. It is known that wound healing and regeneration are physiologically distinct but clearly related processes. In the highly regenerative axolotl, it has been shown that the cell layer formed during the wound healing process produces signals necessary for regeneration^4,5^. Since wound healing must necessarily proceed regeneration, it has long been thought that the two processes could not be separated. Confusing the matter further, similar pathways, such as signaling downstream of reactive oxygen species (ROS), are known to regulate both healing and regeneration. But how the same ROS-mediated signaling can regulate both processes at the same wound site in certain animals is not fully understood.

ROS are highly reactive, oxygen containing byproducts of cellular metabolism. Initially identified as inducers of oxidative stress and cell damage^6^, ROS are now also recognized as positive regulators of cell signaling, stem cell states, host defense, and injury signaling^7^. Biologically relevant ROS include: superoxide (O _2_ ^-^), the hydroxyl radical (·OH), and hydrogen peroxide (H_2_O_2_); but H_2_O_2_ is often considered the major signaling molecule and can act as a second messenger^8^. Additionally, ROS can directly regulate downstream protein activity post-translationally by interacting with thiol groups via redox signaling. Specifically, cysteine residues (which exist as a thiolate anion, Cys-S^-^) can be oxidized by ROS to Cys-SOH at physiological pH, which causes allosteric changes to the protein and alters function^9,10^.

Importantly, the production of injury-induced ROS is a conserved wound response in both plants and animals^11^. The data reveal that ROS accumulation peaks during both wound healing and regeneration, and ROS inhibition has been shown to both delay wound closure rates and prevent regeneration in multiple animal models^12-17^. ROS are capable of inducing vasoconstriction and the recruitment of leukocytes to the wound site after injury^18,19^. In addition, ROS aid in pathogen defense (such as when they are released by phagocytes) and in tissue repair (such as when they induce the proliferation and migration of keratinocytes)^20,21^. In animals capable of regeneration, ROS accumulation has been shown to be sustained after the wound healing process and is required for regeneration to either be initiated or proceed^12-14^.

Responses to ROS evolutionarily range from the activation of transcription factors in prokaryotes to promoting cellular responses (such as cell migration and proliferation) in humans^22,23^. Even cellular outcomes within the same organism can vary depending on ROS levels. Too much ROS can result in cell damage and cell death (or apoptosis), while moderate levels are required to maintain cellular homeostasis or initiate different cell signaling pathways depending on concentration and context^24^. For therapeutic manipulation, understanding this delicate balance of ROS levels will be critical. For example, excess ROS is a leading cause of chronic wounds^25^, while ROS misregulation has been linked to numerous diseases including cancer, Parkinson’s, obesity, diabetes, and cardiovascular disease^26,27^. In addition to concentration, the timing of ROS accumulation is equally important. Activation of ROS-reducing antioxidants is a hallmark of many cancers (since increased ROS can lead to apoptosis, preventing tumorigenesis), yet overproduction of ROS at the right time promotes tumor progression^26,28^. Thus, it will be essential to identify the differences in the amount and timing of ROS that switch downstream signaling from control of wound healing to regrowth.

The aim of this study was to understand the separate role(s) of ROS signaling in wound healing versus regeneration in the same animal model. We chose early injury signaling in planarian flatworms as our model system, as planarians not only possess virtually unlimited regenerative potential (being able to replace all tissues in the organism) but also have a well-characterized wound healing response. Planarian wound healing begins with rapid muscle contractions at the wound site to minimize surface area, followed by mucus secretion from rhabdites over the wound surface^29^. A disorganized epithelial covering is then formed over the wound by the stretching and elongation of cells adjacent to the wound^29-31^. A generic wound response involving an early transcriptional wave occurs and is accompanied by both wound site-specific apoptosis from ∼1-4 hours and body-wide proliferation of adult stem cells from ∼2-6 hours^32-34^. At ∼6 hours, there is increased contact between the wound border and wound tissue itself, as well as an organized appearance of the migrating epithelial cells^29,35^. Lastly, proliferation and migration of new cells to the wound site fully repairs the injury in a more permanent form^29^. The generic response takes place even if regeneration is not initiated by a second injury-specific transcriptional wave^36^. During regeneration, a second peak of adult stem cell proliferation occurs at ∼48-72 hours that results in the formation of an undifferentiated mass at the wound site called the blastema; body-wide apoptosis associated with tissue remodeling follows starting at 3 days^33,34^. Together, the tissue remodeling and differentiation of blastemal cells combine to replace missing structures^37^.

In planarians, ROS have been shown to be required for proper brain patterning, stem cell proliferation, and activation of mitogen activated protein kinase (MAPK) signaling during regeneration^38-40^. Our own work has previously demonstrated that planarian blastema formation requires ROS signaling: ROS accumulation promotes the expression of the chaperone heat shock protein 70 (*hsp70*) that is required for the stem cell proliferative response^38^. Here we show that ROS are not only required for regeneration but are also required for wound healing in planarians, regulating c-Jun (*jun-1)* expression to promote cytoskeletal movements during wound closure. We also show that different threshold levels of ROS control wound healing-specific and regeneration-specific gene expression, where lower ROS levels promote *jun-1* expression and wound closure, while higher ROS levels promote *hsp70* expression, blastema formation, and regeneration. Together, this study suggests that ROS may be a master regulator of injury response programs and will help researchers understand the connections between wound healing and regeneration.

## Results

### ROS Accumulate at the Wound Site Soon After Injury

To assess the role of ROS during wound healing, we amputated planarians in half (bisection) and observed wound morphology every 15 minutes following injury. Consistent with our previous findings^41^, we observed that most animals (70%; n ≥ 44) close their wounds by one hour post amputation (Fig. 1A,C), where a lack of wound closure was defined as mesenchymal tissue protruding from the wound site. Our previous work revealed that ROS accumulate at the wound site one hour after injury^38^. In order to clarify when ROS initially increase, we visualized ROS accumulation using the general oxidative stress indicator dye 5-(and-6)-chloromethyl-2′,7′-dicholorodihydrofluorescein diacetate (CM-H_2_DCFDA) during the first hour following injury in 15-minute increments. We found that there was a sharp increase in ROS accumulation at the wound site between 30 and 45 minutes post amputation that was sustained through the first hour (Fig. 1B,D).

**Fig. 1.**
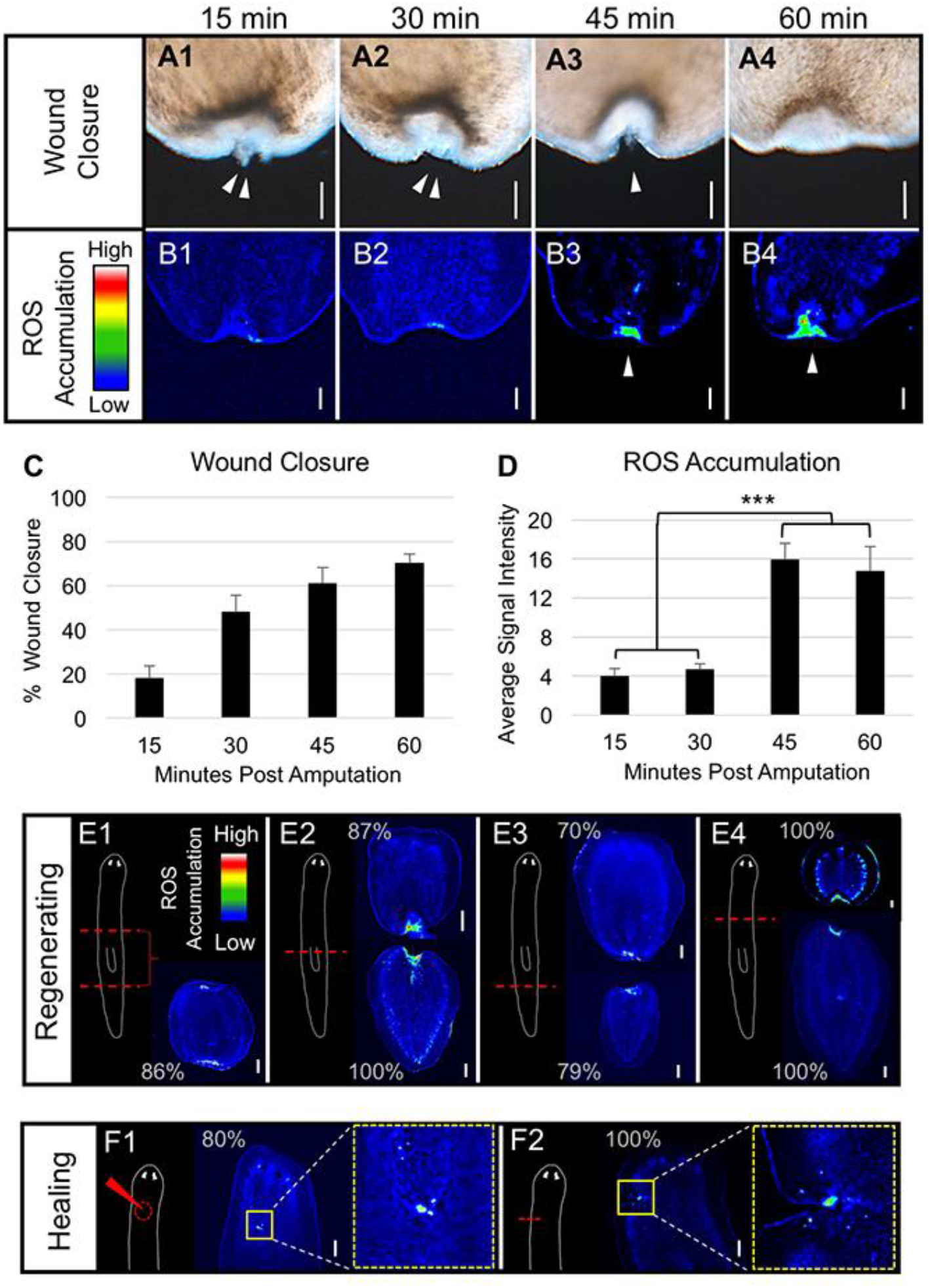
Reactive oxygen species (ROS) accumulate at wound sites regardless of wound type. **(A)** Normal posterior wound closure over the course of one hour post injury. Arrows: presence of mesenchymal tissue emerging from the wound site. Scale bars: 100 μm. (**B**) Posterior ROS accumulation detection during the first hour after injury visualized by the general oxidative stress indicator dye 5-(and-6)-chloromethyl-2′,7′-dicholorodihydrofluorescein diacetate (CM-H_2_DCFDA). Arrows: increased ROS at the wound site. Scale bars: 100 μm. (**C**) Quantification of (A); wound closure over the first hour following amputation. n ≥ 44. Error bars: SEP. (**D**) Quantification of (B); ROS accumulation at posterior wound sites one hour following amputation. n ≥ 15. Error bars: SEM. One-way ANOVA with Tukey’s multiple comparison test: ***P < 0.001. (**E**) ROS accumulation visualized by CM-H_2_DCFDA in various regenerating wounds at one hour (cut diagrams on left of each panel). n ≥ 9. Scale bars: 100 μm. (**F**) ROS accumulation visualized by CM-H_2_DCFDA in various healing wounds at one hour. Cut diagrams to the left; enlargement of boxed injury area on right. n ≥ 10. Scale bars: 200 μm. For all: anterior is up.

For these initial investigations we only examined posterior wounds. However, planarians can survive many different injury types: wounds that remove large amounts of tissue (such as bisection) will heal and regenerate new tissue (the blastema), while those that do not remove tissue will heal but do not regenerate^42^. We examined ROS accumulation one hour post injury in several different injury types. For regenerating wounds, amputations across the anterior-posterior axis regardless of position, as well as regenerates with two wound sites, resulted in an upregulation of ROS at every wound site (Fig. 1E). ROS also accumulated at healing-only wounds induced by either a needle poke or a slice (slit cut) (Fig. 1F), consistent with previous studies ^40^. Together, these data indicate that ROS accumulate during the first hour after injury, before wound closure is complete, regardless of injury type or location.

### ROS are Required for Wound Closure

Previously, we and others have demonstrated that ROS accumulation is required for new tissue growth during blastema formation in planarians^38,39^, consistent with its known role in regeneration in other systems^12-14,16^. Even more so than regeneration, ROS are well established regulators of the wound healing process^22^. To examine the role of ROS during planarian wound healing, we used the flavoenzyme inhibitor diphenyleneiodonium (DPI) to inhibit ROS accumulation (Fig. 2). We found that DPI concentrations as low as 5 μM and up to 50 μM inhibited wound closure (Fig. 2C), although at higher concentrations, animals were increasingly immobile—a sign of toxicity (see our previous work^41^ for examples). We chose 15 μM DPI as our working concentration, as this dose blocked wound healing without affecting normal movement. Our data showed that ROS inhibition in bisected animals significantly prevented wound closure at 45 and 60 minutes (Fig. 2E,F). These are the same time points when the greatest upregulation of ROS at the wound site occurs (Fig. 1B,D). These results suggest ROS are necessary for the wound healing process.

**Fig. 2.**
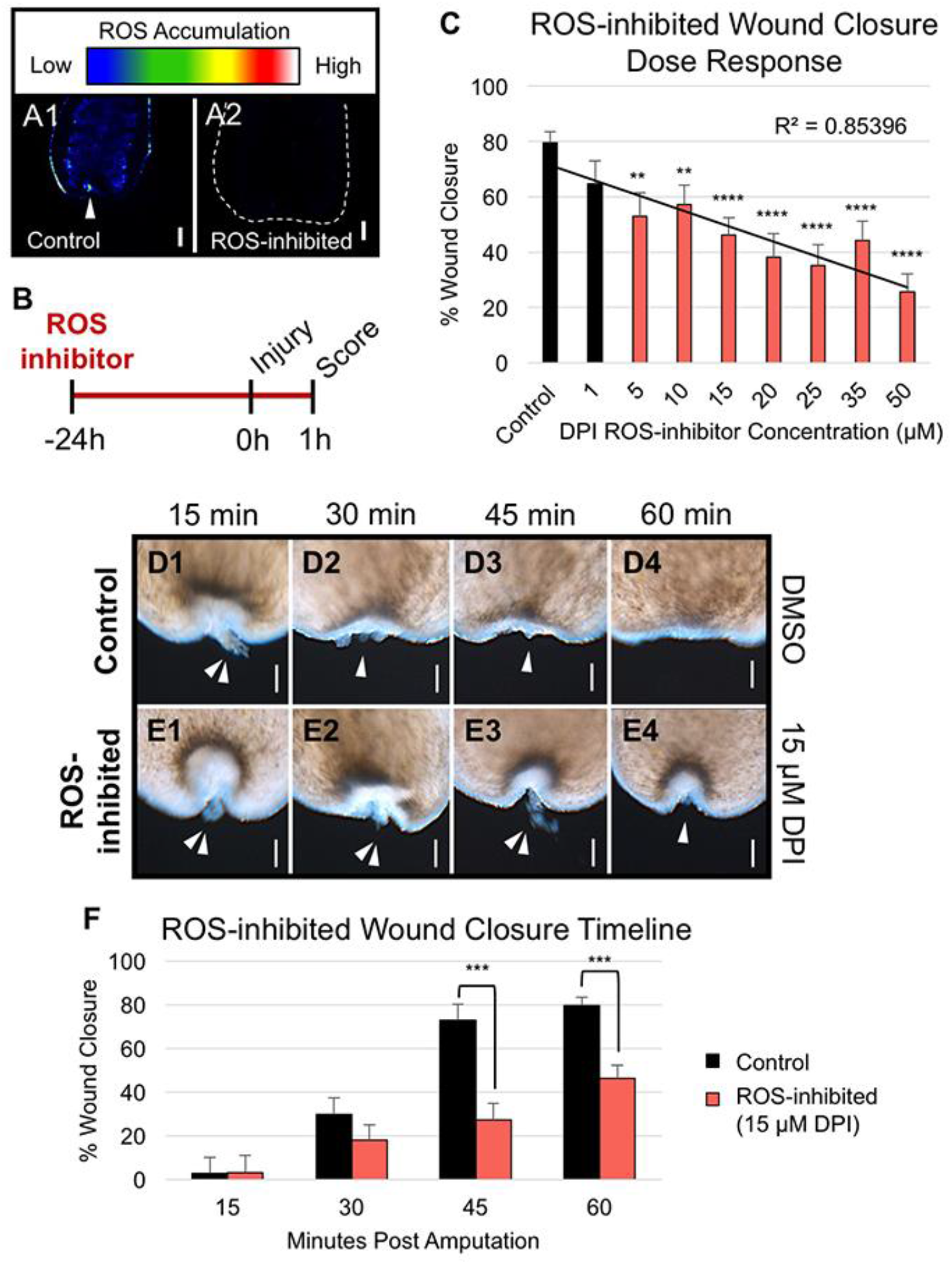
ROS is Required for Planarian Wound Closure. **(A)** ROS accumulation visualized by CM-H_2_DCFDA following treatment with dimethyl sulfoxide (DMSO) for controls (n = 4) or 15 μM diphenyleneiodonium chloride (DPI) for ROS-inhibited (n = 5). Dotted line: outline of ROS-inhibited animal. Arrow: increased ROS at the wound site. Scale bars: 50 μm. (**B**) Experimental timeline for ROS-inhibited wound healing in panels (C-F). Planarians were pre-treated for 24 hours and during wound closure with 15 μM DPI for ROS-inhibited (or DMSO for controls). **(C**) Posterior wound closure after one hour following treatment with increasing DPI concentrations. n ≥ 34. Error bars: SEP. Two sample *t*-test between percents against controls: ** P ≤ 0.01; **** P ≤ 0.0001. (**D**) Control wound closure over one hour. Arrows: presence of mesenchymal tissue emerging from the wound. Scale bars: 50 μm. (**E**) ROS-inhibited wound closure over one hour. Arrows: presence of mesenchymal tissue emerging from the wound. Scale bars: 50 μm. (**F**) Quantification of (D-E); control and ROS-inhibited wound closure over the first hour following amputation. n ≥ 33. Error bars: SEP. Two sample *t*-test between percents against controls: *** P ≤ 0.001. For all: anterior is up.

To validate the results of our pharmacological inhibition of ROS, we next sought to rescue wound closure with exogenous ROS (Fig. 3). H_2_O_2_ is one of the most important ROS signaling molecules in redox biology, which at homeostatic levels is able to initiate pro-survival gene regulation^43^. We first used a dose response to understand the effects of exogenous H_2_O_2_ on wound closure. While concentrations below 300 μM had no effect, higher concentrations of H_2_O_2_ significantly reduced the rate of wound closure (Fig. 3B), presumably due to increased apoptosis associated with higher H_2_O_2_ levels^43^. Therefore, we chose 300 μM H_2_O_2_ to attempt a rescue of wound closure in ROS-inhibited (DPI treated) animals following bisection. We found that DPI-exposed animals subsequently treated with H_2_O_2_ upon injury regained control levels of wound closure compared to DPI exposure alone (Fig. 3D,E). These data highlight a role for ROS in regulating wound healing in planarians and suggest that signaling downstream of ROS is required for wound closure.

**Fig. 3.**
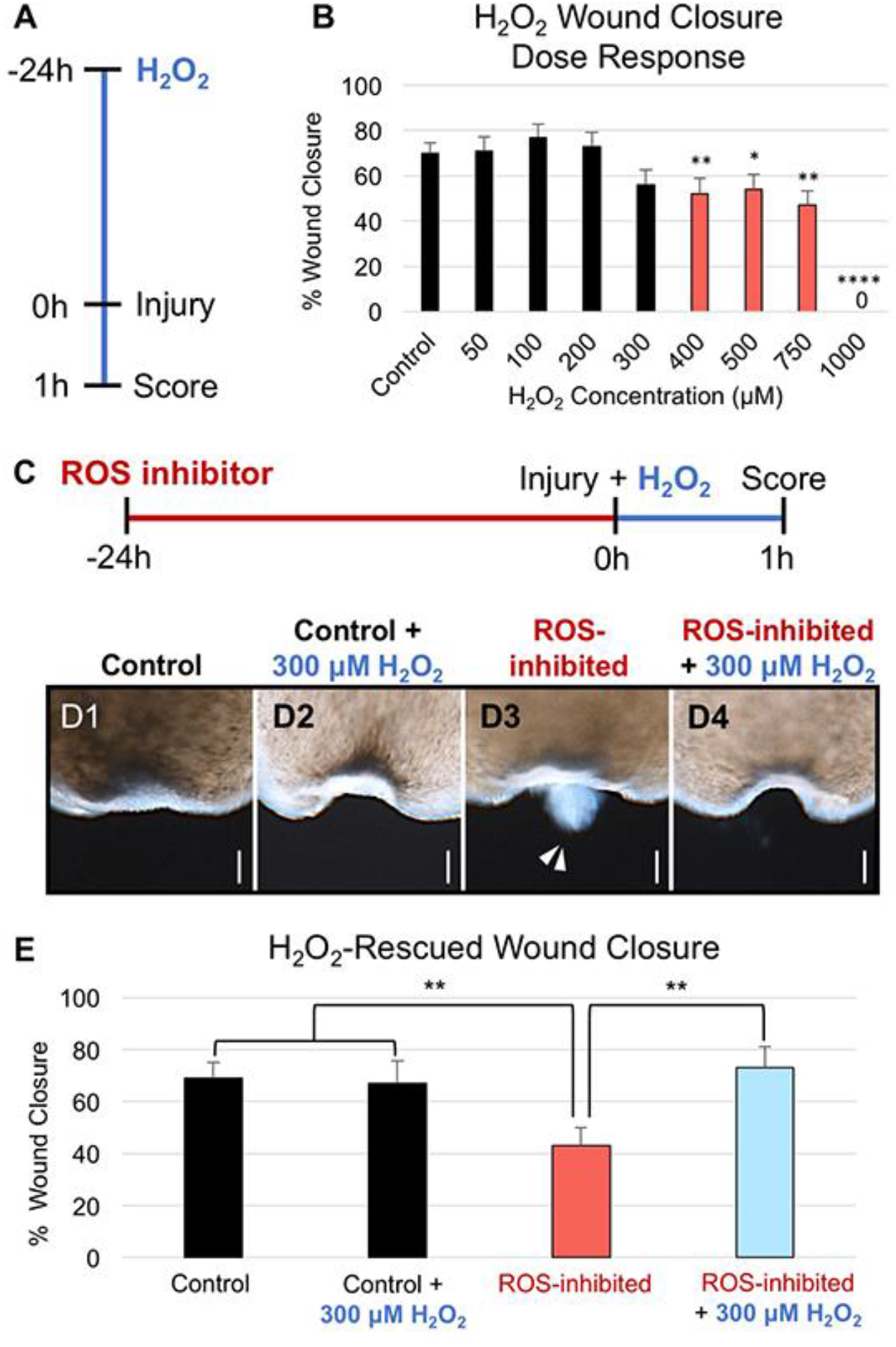
ROS-inhibited Wound Closure is Rescued by Exogenous H_2_O_2_. (**A**) Experimental timeline for H_2_O_2_-exposed wound closure in panel (B). Planarians were pre-treated for 24 hours and during wound closure with H_2_O_2_. (**B**) Posterior wound closure at one hour post injury following treatment with increasing H_2_O_2_ concentrations. n ≥ 52. Error bars: SEP. Two sample *t*-test between percents against untreated controls: * P ≤ 0.05; ** P ≤ 0.01; **** P ≤ 0.0001. (**C**) Experimental timeline for H_2_O_2_-rescued wound closure in panels (D-E). Planarians were pre-treated for 24 hours with 15 μM DPI (red), and then during wound closure with 300 μM H_2_O_2_ (blue). (**D**) ROS-inhibited wound closure rescued by H_2_O_2_ exposure at the time of injury. Arrows: presence of mesenchymal tissue emerging from the wound. Scale bars: 50 μm. Anterior is up. (**E**) Quantification of (D); H_2_O_2_-rescued wound closure. n ≥ 30. Error bars: SEP. Two sample *t*-test between percents against respective controls: ** P ≤ 0.01.

### Injury-Induced Muscle Contractions Do Not Require ROS

Wound healing is a multicomponent process that begins with wound closure. First, muscle contractions help to minimize wound surface area^29^. These contractions result in a dark ring of pigment surrounding the wound (Fig. 2D1) that lessens as the muscles relax with the formation of the initial epithelial covering (Fig. 2D4). We found that ROS loss did not affect these contractions. Similar to controls, ROS-inhibited wounds had pigment rings immediately after injury at 15 minutes (Fig. 2E1), suggesting muscle contraction does not require ROS. However, unlike control wounds, contraction persisted in ROS-inhibited wounds, marked by the retention of the dark pigment ring at one hour post injury (Fig. 2E4) concurrent with failure to close the wound. Following muscle contractions, normal wound healing requires epithelial cells adjacent to the wound undergo cytoskeletal changes to provide an initial covering^30^. A study using magnesium chloride to inhibit planarian muscle contraction during wound closure still observed this epithelial cell spreading^44^, suggesting muscle contraction and cytoskeletal movements may use independent mechanisms. Together with our findings, these data suggest that ROS are not required for muscle contraction during wound healing.

### ROS Regulate Actin-Mediated Epithelial Stretching and Reorganization

Following contraction, the initial epithelial covering forms by actin-mediated cytoskeletal changes, including the elongation (stretching) and rearrangement of adjacent cells over the wound surface^45,46^. To investigate whether ROS play a role in this step of wound closure, we chose to visualize filamentous actin (F-actin) using phalloidin labeled with fluorescein isothiocyanate (FITC), in combination with Hoechst staining of nuclei to distinguish individual cells (Fig. 4). When cells are labeled in this manner, the intact (uninjured) planarian epidermis appears honeycombed in appearance, outlining the tight network of phalloidin-FITC-labeled cell borders (green) surrounding cell nuclei (blue) and the openings (black) for mucosal secretions by rhabdomeres^30^ (Fig. 4A1). A side view of the same epidermis (Fig. 4B) reveals the regular dorsal alignment of cell nuclei in the single cell layer that comprises the planarian epidermis (a representative trace drawing of the dorsal epidermal shape, as seen from the side, is provided to the left of the panel). To examine the epithelium during wound healing, injury to the dorsal anterior region using a hypodermic needle allowed for easy observation of wound closure and the wound margins (Fig. 4A2). Side views of the wound (Fig. 4C) revealed that during the first hour after injury, the missing tissue at the wound site (black region in right panel of Fig. 4C1) was rapidly replaced by actin-labeled structures/cells (Fig. 4C2-C4). By 60 minutes post-injury, wounds regained the smooth, uninterrupted dorsal covering seen in the intact epidermis (Fig. 4B). However, healing wounds now possessed actin staining where the wound opening had previously been (Fig. 4C4), suggesting that cells stretched to form the wound epithelium.

**Fig. 4.**
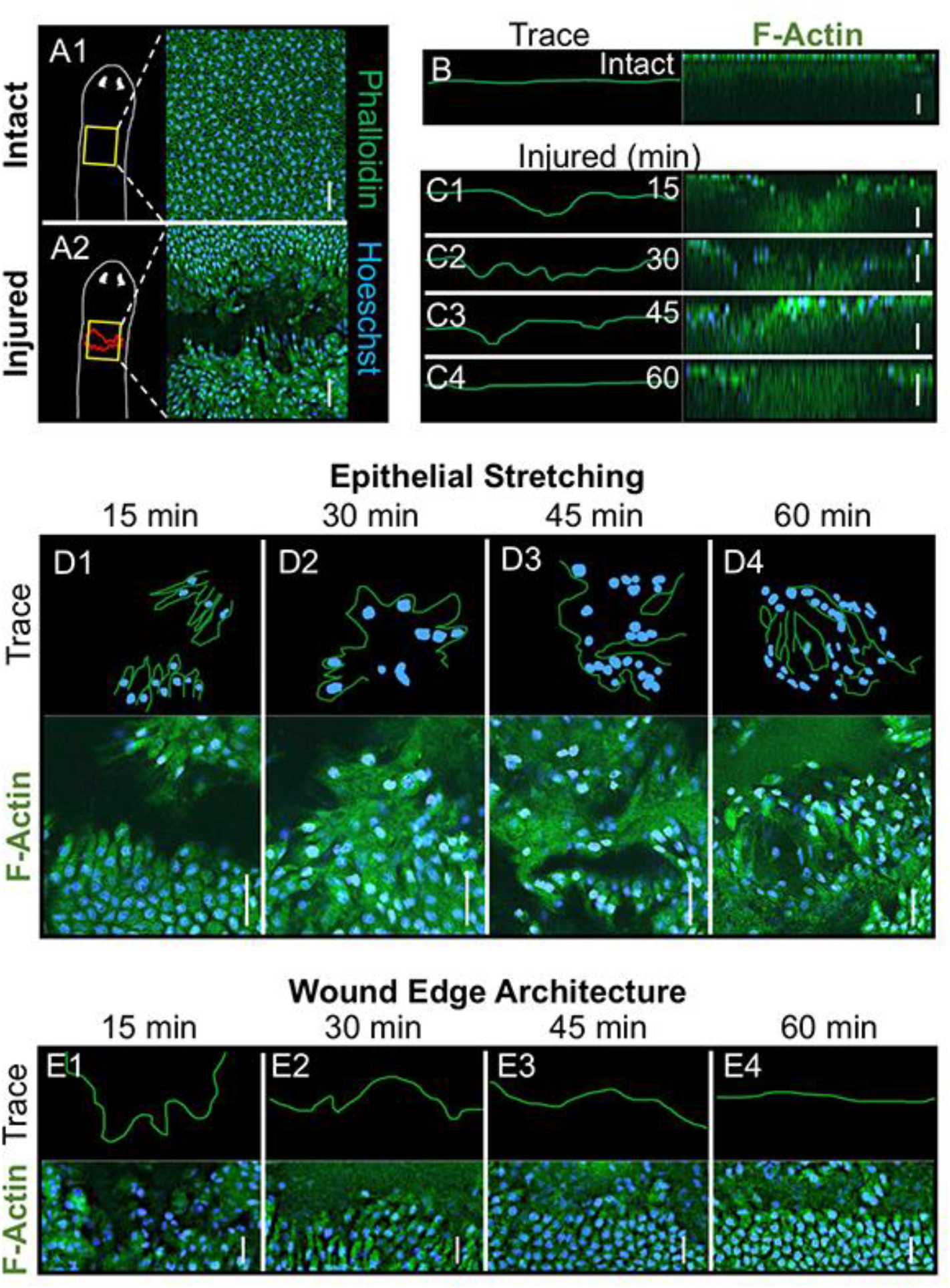
Wound Closure is Actin-Mediated. F-actin (green, Phalloidin-FITC) and nuclei (blue, Hoechst 33342) labeling. (**A**) F-actin and nuclei labeling of dorsal anterior epidermis in intact versus injured animals at one hour post injury (dorsal needle punch). Scale bars: 25 μm. Anterior is up. Dorsal faces reader. (**B**) Left panel: representative trace of dorsal epidermis shape from image in right panel. Right panel: XZ image of confocal Z-stack of an intact planarian with F-actin and nuclei labeling. Scale bar: 25 μm. Dorsal is up. (**C**) Wound landscape every 15 minutes during the first hour following injury as depicted in (A2). Left panels: representative traces of dorsal epidermis shape. Right panels: XZ images of confocal Z-stacks of injured planarians with F-actin and nuclei labeling. Scale bars: 25 μm. Dorsal is up. (**D**) Epithelial elongation (stretching) of wound edge throughout the first hour after wounding as in (A2). Top panels: representative traces of stretching structures from each time point. Bottom panels: F-actin and nuclei labeling showing epithelial stretching over time. Scale bars: 25 μm. Dorsal view. (**E**) Wound edge architecture over the first hour after wounding as in (A2). Top panels: representative traces of edge appearance. Bottom panels: F-actin and nuclei labeling showing edge organization. Scale bars: 15 μm. Dorsal view. For all conditions: n = 7.

To further understand the actin-cytoskeletal changes occurring during wound closure, we examined other focal planes of the wound site. Dorsal views of the wound every 15 minutes post-injury allowed for analyses of wound-site epithelial stretching (Fig. 4D). Over the first hour, cells on either side of the wound (Fig. 4D1) elongated and moved to cover the opening (Fig. 4D2-D4), as epithelial stretching increased in complexity (representative trace drawings of cell shape and location changes are provided above the images). These cell rearrangements formed what appeared to be scaffolding for cells to repair the site of injury (Fig. 4D4). Focusing the dorsal view on the wound margins allowed for analyses of wound edge architecture (Fig. 4E). We observed that over the first hour after injury, the organization of wound edges went from highly disorganized (Fig. 4E1) to containing regions of straight alignment that bordered the wound (Fig. 4E2). Collectively, this combination of actin rearrangements resulted in the smooth initial epithelial covering as seen in Fig. 1A4.

Our data revealed that ROS are required for normal wound closure in planarians (Fig. 2). ROS promote wound healing in part by regulating actin-mediated wound closure^47^. Therefore, we hypothesized that ROS signaling promotes cytoskeletal movements during wound closure in planarians. We found that, similar to bisected animals, after dorsal wounding ROS-inhibited (DPI treated) animals also failed to close their wounds by one hour post-injury (Fig. 5A). This was concurrent with a lack of actin-labeled cell structures covering the entire wound site (open arrow/black gaps in Fig. 5B and uneven nuclear trace Fig. 5C2). Furthermore, ROS inhibition prevented epithelial stretching (Fig. 5D), as well as cell reorganization at the wound edge (Fig. 5E). Interestingly, ROS-inhibited wounds at one hour (Fig. 5) more closely compared to control wounds at 0-15 minutes (Fig. 4) rather than time-matched controls. Together, these results demonstrate that ROS are required for the actin-mediated epithelial cell rearrangements that drive wound closure.

**Fig. 5.**
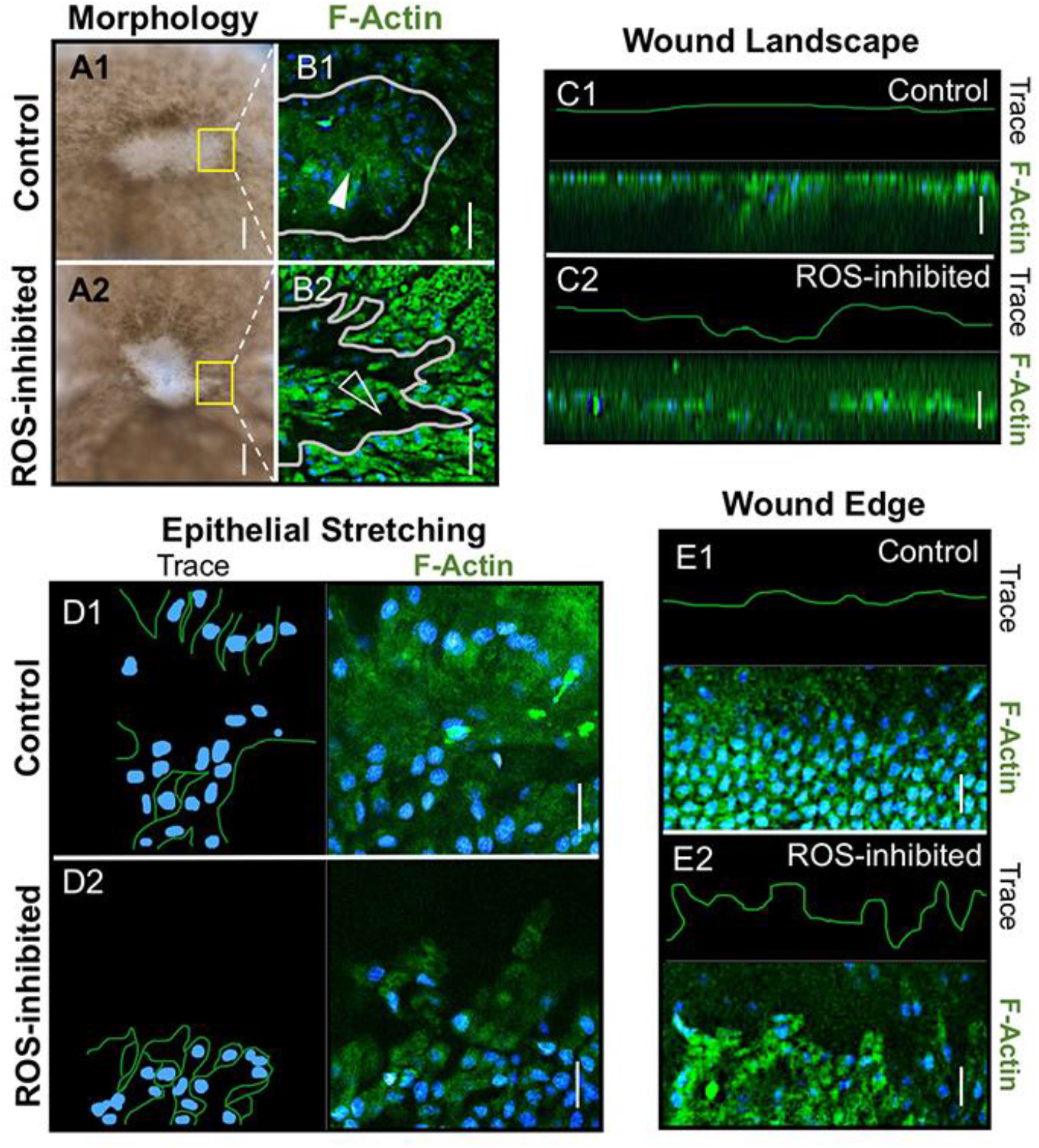
ROS Regulates Epithelial Stretching and Organization During Wound Closure. Control (DMSO) and ROS-inhibited (15 μM DPI) dorsal punch wounding (as depicted in Fig. 4A2) with F-actin (green, Phalloidin-FITC) and nuclei (blue, Hoechst 33342) labeling. (**A**) Wound morphology at one hour post injury in control and ROS-inhibited wounds. 5/7 ROS-inhibited versus 2/7 control wounds remained open. Scale bars: 100 μm. Dorsal view. (**B**) F-actin and nuclei labeling of wound edges in control and ROS-inhibited wounds at one hour. Images correspond to area in yellow boxes in (A). 1/7 ROS-inhibited versus 9/9 control wounds had actin cover the entire wound site. Solid grey lines: wound edges. Solid arrow: presence of actin staining in wound site. Open arrow: gap in actin staining. Scale bars: 25 μm. Dorsal view. (**C**) Wound landscape at one hour in control and ROS-inhibited injuries. Top panels: representative traces of dorsal epidermis shape. Bottom panels: XZ images of confocal Z-stacks of injured planarians with F-actin and nuclei labeling. 7/7 ROS-inhibited versus 1/9 control wounds displayed disorganized dorsal landscape. Scale bars: 25 μm. Dorsal is up. (**D**) Control and ROS-inhibited epithelial elongation (stretching) at one hour. Left panels: representative traces of stretching structures. Right panels: F-actin and nuclei labeling. 0/7 ROS-inhibited versus 8/8 control wounds displayed complex epithelial stretching. Scale bars: 15 μm. Dorsal view. (**E**) Wound edge architecture in control and ROS-inhibited wounds at one hour. Top panels: representative traces of edge appearance. Bottom panels: F-actin and nuclei labeling showing edge organization. 0/7 ROS-inhibited versus 9/9 control wounds exhibited organized wound edges. Scale bars: 15 μm. Dorsal view.

### *ROS Signaling is Required for Wound Site* jun-1 *Expression*

In planarians, the c-Jun homolog *jun-1* is part of a generic wound response that includes a wave of gene expression upregulated as early as 30 minutes that is specifically expressed in epidermal cells^32,36^. Since it was also found to promote wound healing in other animals, we hypothesized that *jun-1* expression might be regulated by ROS signaling during wound closure. We found that, similar to ROS accumulation, *jun-1* expression was significantly upregulated at the wound site, with highest wound site expression between 45 minutes and 1 hour (Fig. 6A). Consistent with a role in early wound healing events (such as wound closure), our data revealed that wound site *jun-1* expression decreases after two hours (Fig. 6A5). To investigate whether ROS accumulation is required for injury-induced *jun-1* expression, we examined ROS-inhibited (DPI treated) animals at 45 minutes after bisection (Fig. 6B). Our results showed that ROS inhibition significantly decreased *jun-1* expression compared to controls (Fig. 6B,C). These data suggest that ROS accumulation is required for *jun-1* expression at the wound site, raising the possibility that *jun-1* transduces ROS signals to control wound closure during wound healing.

**Fig. 6.**
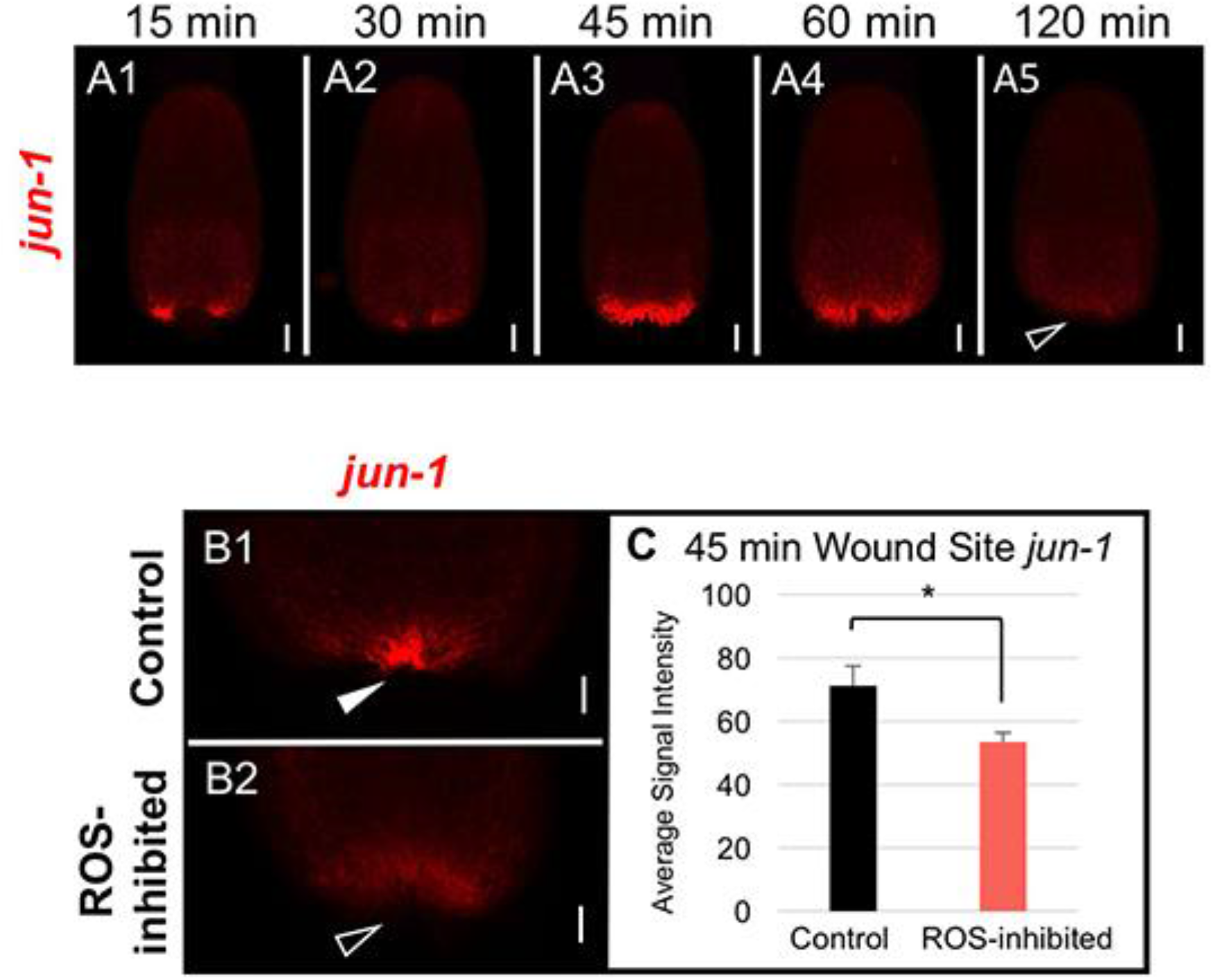
ROS Regulates *jun-1* Expression at the Wound Site. Expression of the transcription factor c-Jun *(jun-1)* visualized by fluorescent *in situ* hybridization (FISH). (**A**) Posterior wound *jun-1* expression in bisected planarians after amputation. Open arrow: absence of wound site expression. Scale bars: 100 μm. (**B**) Control (DMSO) and ROS-inhibited (15 μM DPI) posterior wound *jun-1* expression at 45 minutes post injury in bisected fragments (using treatment scheme as depicted in Fig. 2B). Solid arrow: presence of upregulated wound site gene expression. Open arrow: inhibition of wound site expression. Scale bars: 200 μm. (**C**) Quantification of (B); control and ROS-inhibited wound site *jun-1 expression* at 45 minutes. Error bars: SEM. Student’s *t*-test: * P ≤ 0.05. For all: n = 10; anterior is up.

### *Epithelial Stretching, but not Wound Edge Reorganization, is Mediated by* jun-1

c-Jun also has been shown to play an important role in promoting epithelial cell migration^48^. Therefore, we next investigated whether *jun-1* was required for ROS-mediated wound-related epithelial movements in planarians. We used RNA interference (RNAi) to knockdown *jun-1* expression (Fig. 7A) and then examined its effects on wound closure. We found that dorsal wounding (as in Fig. 4A2) of *jun-1* RNAi animals produced wounds similar to ROS-inhibited wounds, in that they both had a disorganized epidermis that lacked actin-labeled structures filling the wound site (Fig. 7B) concurrent with inhibited epithelial stretching (Fig. 7C). Phenotypically, both *jun-1* RNAi and ROS-inhibited wounds at one hour post injury more closely resembled control wounds at 15 minutes (Fig. 4) rather than time-matched controls, suggesting that wound closure events had stalled. However, DPI treated wounds differed from *jun-1* RNAi wounds, as loss of *jun-1* did not affect wound edge reorganization (Fig. 7D). This was unlike the effects observed after ROS inhibition: while *jun-1* RNAi wound edges were evenly aligned similar to controls (Fig. 7D2), DPI treatment resulted in highly disorganized wound edges (Fig. 4E2). These data suggest that during planarian wound closure, *jun-1* regulates actin-mediated epithelial cell stretching but does not regulate cell rearrangements at the wound margin.

**Fig. 7.**
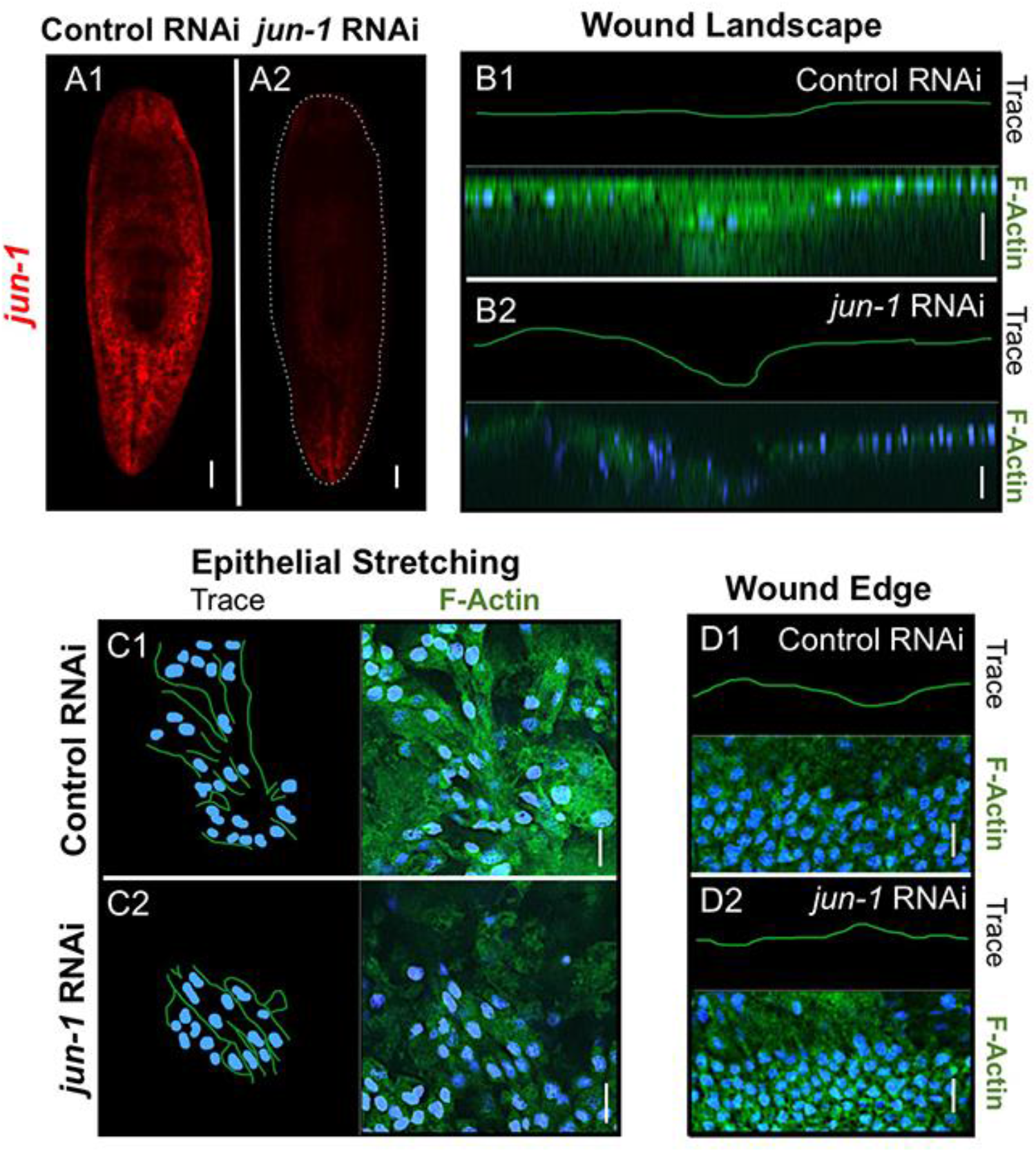
Epithelial Stretching During Wound Closure is Mediated by *jun-1*. Control (Venus-GFP) and *jun-1* RNA interference (RNAi). (**A**) *jun-1* expression in intact animals as visualized by FISH (red). 7/10 *jun-1* RNAi animals versus 1/10 controls showed reduced *jun-1* staining. Dotted line: outline of *jun-1* RNAi animal. Scale bars: 100 μm. Anterior is up. (**B-D**) F-actin (green, Phalloidin-FITC) and nuclei (blue, Hoechst 33342) labeling of dorsal punch wounds (as depicted in Fig. 4A2). **(B)** Wound landscape at one hour post injury in control RNAi and *jun-1* RNAi wounds. Top panels: representative traces of dorsal epidermis shape. Bottom panels: XZ images of confocal Z-stacks of injured planarians with F-actin and nuclei labeling. 6/6 *jun-1* versus 1/7 control wounds displayed a disorganized dorsal landscape. Scale bars: 25 μm. Dorsal is up. (**C**) Control RNAi and *jun-1* RNAi epithelial elongation (stretching) structures at one hour. Left panels: representative traces of stretching structures. Right panels: F-actin and nuclei labeling. 0/7 *jun-1* versus 7/7 control wounds displayed complex epithelial stretching. Scale bars: 15 μm. Dorsal view. (**D**) Wound edge architecture in control RNAi and *jun-1* RNAi wounds at one hour. Top panels: representative traces of edge appearance. Bottom panels: F-actin and nuclei labeling showing edge organization. All ROS-inhibited (7/7) and control wounds (7/7) exhibited organized wound edges. Scale bars: 15 μm. Dorsal view.

### ROS Regulate Distinct Wound Healing and Regeneration Signaling Programs

The data from this study identify ROS as essential regulators of wound healing in planarians, while our previous work demonstrated that ROS are also essential regulators of planarian regeneration^38^. These findings are consistent with observations that ROS accumulation increases during both wound healing and regeneration events, and that inhibition of ROS delays wound closure and prevents regeneration^12-15^. Yet, the relationship between wound healing and regenerative ROS programs remains unclear. Given the related natures of wound healing and regeneration, we hypothesized that either 1) a single ROS signaling cascade sequentially regulated both wound healing and regeneration, or 2) separate ROS signaling pathways controlled each process individually.

To tease apart these opposing hypotheses, we examined the expression of key ROS-mediated genes that were specifically associated with each process (Fig. 8). From this study, we identified *jun-1* as specifically upregulated by ROS during wounding (Fig. 6). We had previously identified the chaperone heat shock protein 70 (*hsp70*) as specifically upregulated by ROS during planarian regeneration, upstream of stem cell-mediated new tissue growth^38^. We also took advantage of the fact that we had injury protocols that were specific to each process. Injuries that do not remove tissue, such as lateral slit cuts (Fig. 8A), heal without regenerating. Larger injuries that remove tissue, such as bisection (Fig. 8B), heal and then regenerate, forming the new tissues of the blastema (white tissues at the solid arrow in Fig. 8B that are absent at the open arrow in Fig. 8A).

**Fig. 8.**
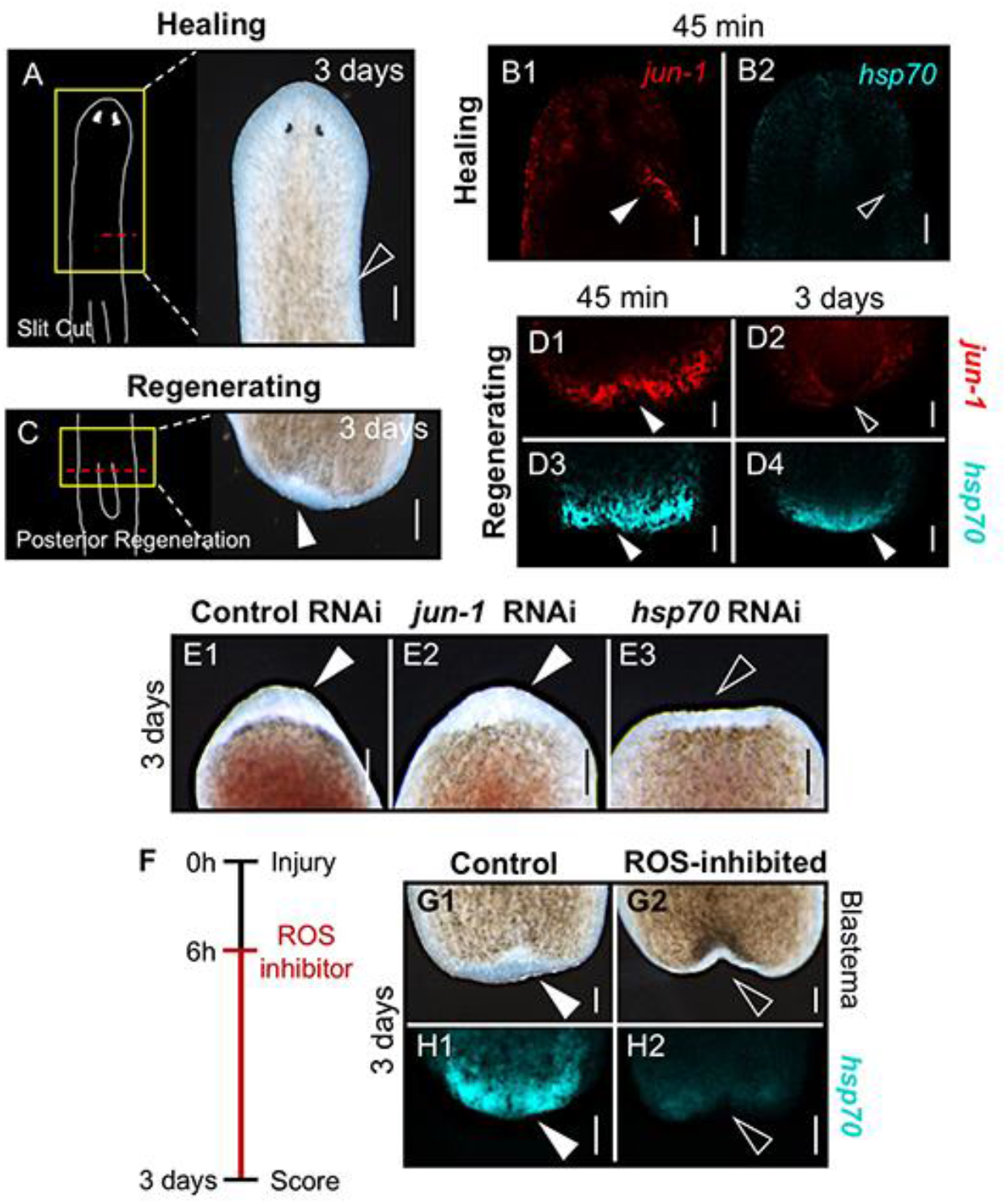
Different ROS-Mediated Gene Expression Distinguishes Wound Healing from Regeneration. (**A**) Healing wound (cut diagram on left). Injuries without tissue removal do not regenerate. Open arrow: lack of blastema formation at 3 days post injury. n = 20. Scale bar: 200 μm. (**B**) Injury site *jun-1 and* Heat shock protein 70 (*hsp70*) expression visualized by FISH at 45 minutes post injury in healing wounds as in (A). Solid arrow: gene expression. Open arrow: absence of gene expression. Scale bars: 100 μm. *jun-1* wound expression compared to uninjured side; n = 11; Student’s *t*-test, P = 0.0164. *hsp70* wound expression compared to uninjured side; n = 12; Student’s *t*-test, P = 0.1085. (**C**) Regenerating wound (cut diagram on left). Solid arrow: blastema at 3 days. n = 21. Scale bar: 200 μm. (**D**) Injury site *jun-1* and *hsp70* expression at 45 minutes and 3 days in regenerating wounds as in (C). Solid arrow: gene expression. Open arrow: absence of gene expression. At 45 minutes: all had both *jun-1* (10/10) and *hsp70* (12/12) expression. At 3 days: 2/10 expressed some *jun-1*, while 10/10 expressed *hsp70*. Scale bars: 100 μm. (**E**) Blastema formation at 3 days following control (Venus-GFP), *jun-1*, and *hsp70* RNAi. Solid arrows: blastema. Open arrow: inhibition of blastema. n = 10. Scale bars: 200 μm. *jun-1* RNAi blastema sizes were comparable to controls; Student’s *t*-test: P = 0.8624. (**F**) Experimental timeline for panels (G-H); delayed ROS inhibition starting at 6 hours post injury (when wound healing is complete). (**G**) Blastema formation at 3 days following delayed treatment; control (DMSO) and ROS-inhibited (15 μM DPI). Solid arrow: blastema. Open arrow: inhibition of blastema. 0/11 delayed ROS-inhibited fragments versus 10/10 controls formed a blastema. Scale bars: 100 μm. (**H**) *hsp70* expression at 3 days following delayed treatment. Solid arrow: gene expression. Open arrow: absence of wound site expression. For both: n = 9. Scale bars: 50 μm. Delayed ROS inhibition significantly blocked expression compared to controls; Student’s *t*-test: P ≤ 0.0001. For all: anterior is up.

We found that in a healing-only injury at 45 minutes, wound healing-associated *jun-1* was significantly upregulated at the wound site but regeneration-associated *hsp70* was not (Fig. 8B). Conversely, in regenerating injuries both *jun-1* and *hsp70* were upregulated at the wound site at 45 minutes, but only the regeneration-associated *hsp70* was upregulated in the blastema at 3 days (Fig. 8D). Since *jun-1* was only expressed at the wound site during wound closure in both wound types, we used RNAi to examine the effects of *jun-1* inhibition on blastema formation (Fig. 8E). Our results revealed that loss of *jun-1* did not significantly affect blastema size (Fig. 8E2) unlike loss of *hsp70* (Fig. 8E3), which has been demonstrated to reduce blastema size^38^. These findings indicate that, even though both are upregulated by ROS after injury, *jun-1* is wound healing specific while *hsp70* is regeneration specific.

Our data show that ROS signaling is required for both early injury-related events (wound closure, 1 hour) and later events (regeneration, 3 days). In addition, we show that *hsp70* is expressed both early and late despite only being required for later regenerative events, as loss of early *hsp70* expression (via *hsp70* RNAi) has no effect on wound closure (Fig. 8E). Thus, it was not clear whether or not early ROS signaling (associated with wound closure) was sufficient on its own to induce regeneration as would be predicted by the single ROS signaling cascade hypothesis. We therefore performed a delayed ROS inhibition assay, where we investigated whether early ROS signaling would be sufficient to overcome later inhibition. We allowed bisected animals to complete wound healing undisturbed, where the early expression of ROS, *jun-1* and *hsp70* were allowed to occur. Then, at 6 hours post-injury (when wound healing is complete), we placed regenerates into DPI to inhibit ROS solely during blastema formation (from 6-72 hours). We found that none of the animals with delayed ROS inhibition formed a blastema (Fig. 8G), coincident with a loss of wound site *hsp70* expression at 3 days (Fig. 8H).

These data do not support our first hypothesis that a single ROS signaling cascade sequentially regulates both wound healing and regeneration. Instead, the data are consistent with our second hypothesis: that separate ROS signaling pathways control each process individually. Our data indicate that strong wound site *hsp70* expression soon after injury signifies that regeneration will occur, even though its early expression alone is not sufficient to promote blastema formation. This is consistent with reports that *hsp70* expression is required for the initiation and maintenance of both muscle and liver regeneration in mice^49,50^. Together, our data suggest that ROS accumulation is upstream of two distinct signaling pathways that regulate wounding healing and regeneration independently.

### Different Threshold Levels of ROS Promote Wound Healing Versus Regeneration

The identification of separate ROS signaling mechanisms regulating wound healing and regeneration still does not provide a mechanistic explanation for why ROS accumulation can promote different downstream gene expression and cellular activities in the same wound. Specifically, in regenerating wounds ROS promote *jun-1*-mediated wound closure and *hsp70*-mediated stem cell proliferation, while in healing wounds ROS promote only *jun-1*-mediated wound closure (Fig. 8). Furthermore, it has been shown that planarians still initiate early (∼4 hours) proliferation and apoptosis following a healing (non-regenerative) injury, and the response is proportional to injury size^33,34^. Therefore, even though ROS accumulate at all wound sites (Fig. 1), we wondered whether the level of ROS accumulation varied by injury type. We examined ROS levels in different types of healing wounds (slit cuts and dorsal punch) and regenerative wounds (anterior and posterior regeneration) (Fig. 9A). Our data reveal that healing wound sites have significantly lower concentrations of ROS accumulation than regenerating wound sites, even when correcting for differences in wound sizes (Fig. 9B).

**Fig. 9.**
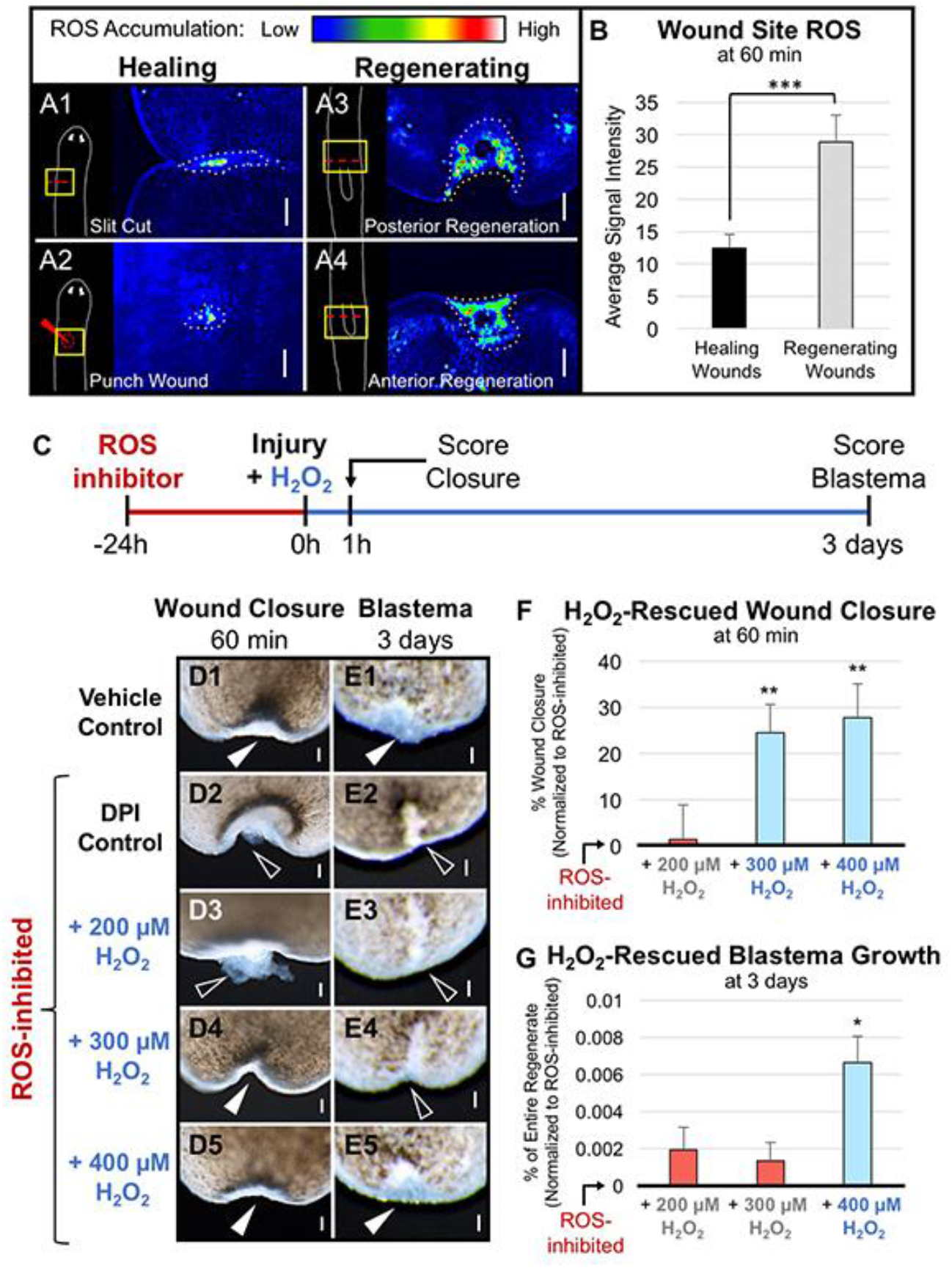
Different ROS Threshold Levels Regulate Wound Healing and Regeneration. (**A**) Wound site ROS accumulation visualized by CM-H_2_DCFDA in healing versus regenerating wounds one hour after injury. Dotted lines: the measured wound region for each injury type. n ≥ 22. Scale bars: 100 μm. (**B**) Quantification of (A); wound site ROS levels in healing (A1-A2) versus regenerating (A3-A4) wounds. Error bars: SEM. Student’s *t*-test: *** P ≤ 0.001. (**C**) Experimental timeline for panels (D-G); H_2_O_2_ rescue of ROS inhibition. Animals were pre-exposed to 15 μM DPI for 24 hours prior to injury, then immediately following injury to H_2_O_2._Exceptions: Vehicle Controls (only pre-exposed to DMSO) and ROS-inhibited DPI Controls (only pre-exposed to DPI). (**D**) ROS-inhibited wound closure with increasing concentrations of H_2_O_2_. Solid arrows: closed wounds. Open arrows: absence of wound closure. n ≥ 37. Scale bars: 50 μm. (**E**) ROS-inhibited blastema growth with increasing concentrations of H_2_O_2_. n ≥ 12. Scale bars: 50 μm. (**F**) Quantification of (D); percent ROS-inhibited wound closure with increasing H_2_O_2_ concentrations, normalized to ROS-inhibited DPI Control levels. n ≥ 37. Error bars: SEP. Two sample *t*-test between percents against ROS-inhibited: ** P ≤ 0.01. (**G**) Quantification of (E); ROS-inhibited blastema growth (relative to body size) with increasing H_2_O_2_ concentrations, normalized to ROS-inhibited DPI Control levels. Error bars: SEM. One-way ANOVA with Tukey’s multiple comparison test: * P ≤ 0.05 versus ROS-inhibited animals. For all: anterior is up.

These data suggest that different relative threshold levels of ROS promote wound healing versus regeneration. This is consistent with the known cellular responses to different ROS thresholds; basal levels of ROS are required to maintain cellular homeostasis and promote cell signaling, while both too little and too much are harmful^51^. We hypothesized that within this physiological (beneficial) range of ROS levels, two separate (sub)thresholds exist: a comparatively lower threshold that initiates wound healing responses, and a comparatively higher level that is required to initiate regeneration (Fig. 10A). In partial support of our hypothesis, we noticed that the optimal concentration of DPI required to inhibit regeneration (10 μM^38^) was lower than the optimal concentration required to inhibit wound closure (15 μM, Fig. 2E), suggesting that regeneration was more sensitive to reductions in ROS levels.

**Fig. 10.**
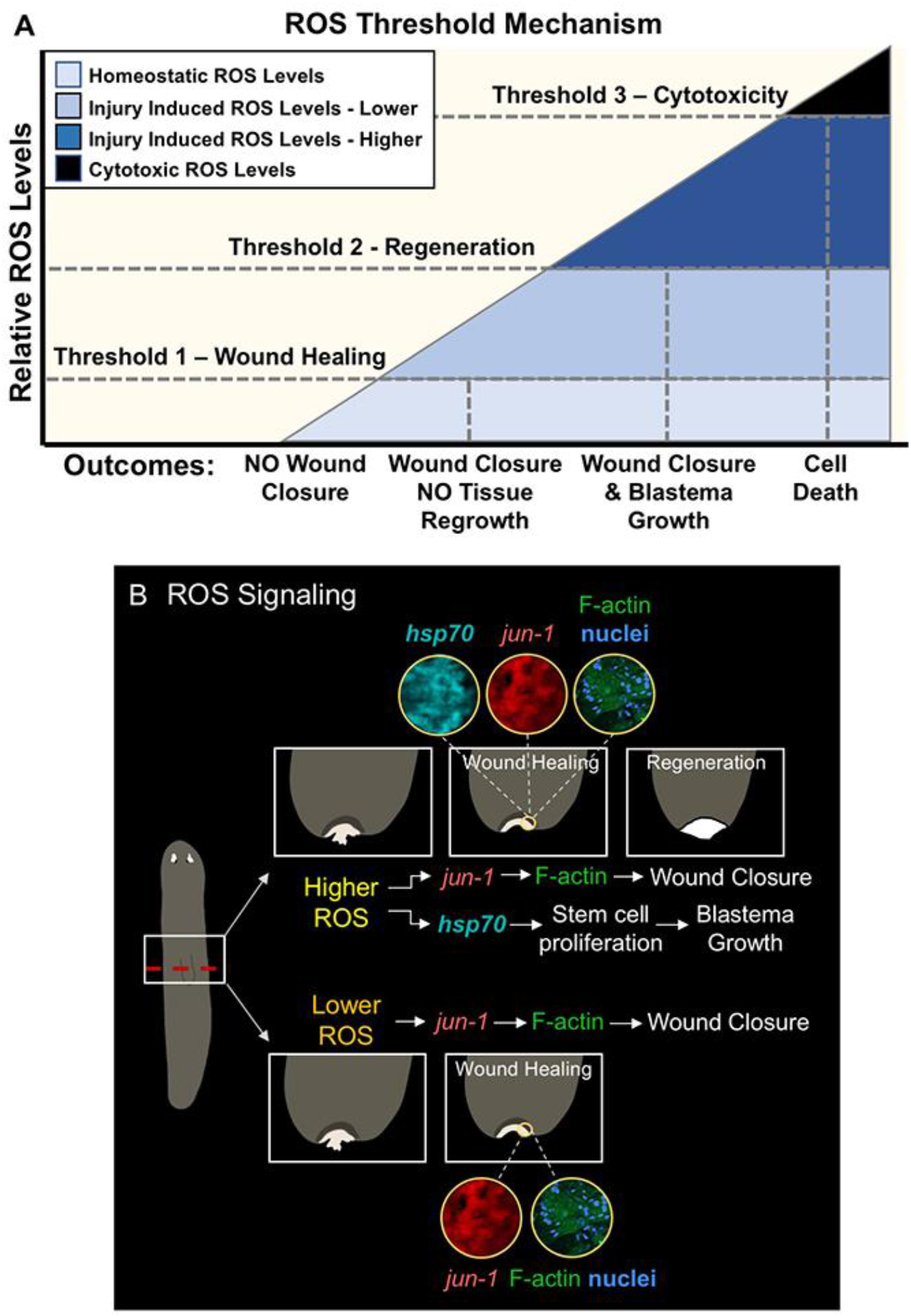
Model of ROS-Mediated Signaling During Tissue Repair. (**A) Diagram of proposed ROS threshold mechanism during repair**. Basal or extremely low ROS levels are not sufficient to promote wound closure. Injuries result in increased ROS accumulation at the wound site, which is required for tissue repair. Within this increased ROS, the data indicate two separate thresholds exist following injury: a lower relative threshold that promotes wound healing and a higher relative threshold that promotes new tissue regrowth, blastema formation, and regeneration. The data also indicate that even higher ROS levels constitute a third threshold to promote oxidative stress, cell damage and death. (**B**) **Model of ROS signaling during repair**. Higher levels of injury-induced ROS in regenerating wounds are sufficient to induce both *jun-1* expression (which drives actin-mediated wound closure) and *hsp70* expression (which drives stem cell proliferation and new tissue growth). Lower levels of injury-induced ROS found in healing wounds are only sufficient to induce *jun-1* expression, leading to wound closure without blastema formation.

To test our hypothesis, we used exogenous ROS exposure to attempt to rescue either wound healing alone or both wound healing and regeneration in ROS-inhibited injuries. Bisected animals were exposed to DPI prior to injury, after which they were treated with different concentrations of H_2_O_2_ (Fig. 9C-G). Our earlier studies (Fig. 3) indicated that 300 μM H_2_O_2_ was sufficient to rescue wound closure in DPI-treated animals, therefore we chose to examine a range centered around that concentration. We found that non-inhibited control animals both closed their wounds at 1 hour and formed a blastema (regenerated) by 3 days as expected, while DPI-treated animals did neither (Fig. 9D,E). Our data revealed that: 200 μM H_2_O_2_ was not sufficient to recuse either wound healing or regeneration following ROS inhibition; 300 μM H_2_O_2_ rescued wound healing but not blastema formation (regeneration); while 400 μM H_2_O_2_ was able to rescue both wound healing and regeneration (Fig. 9F,G). These data demonstrate that lower levels of ROS accumulation will initiate wound healing, while higher ROS levels are required to initiate regeneration, consist with our threshold model (Fig. 10).

## Discussion

Collectively, our findings provide insights into cytoskeletal-mediated mechanisms of early wound healing in planarians, and our investigation of both wound healing and regeneration in the same model system (at the same wound) has enabled us to parse out differences in two processes that were once thought to be inseparable. Our investigations identified ROS, which are well known regulators of both wound healing and regeneration^12-21^, as differential controllers of wound repair mechanisms. We show that ROS-mediated *jun-1* expression is a regulator of epithelial stretching during wound closure, while ROS signaling upstream of *hsp70* expression governs the regenerative program. This occurred in a threshold-dependent manner, suggesting that careful regulation of ROS levels is critical after injury.

The threshold mechanism (Fig. 10A) helps explain why ROS signaling can activate different programs that drive separate morphological outcomes (tissue repair vs. functional tissue replacement) despite the fact that ROS accumulate after injury at all wound sites regardless of type or size. We found that cellular responses to injury are dependent on differential levels of ROS, which is consistent with ROS functioning as positive drivers of tissue repair and immune recruitment^19-21^. For example, in adult zebrafish, wounds that only require healing exhibit ROS accumulation, while injuries involving new tissue formation require sustained ROS (specifically H_2_O_2_) to promote the regeneration process^12,52^. However, in other instances, ROS are known to promote fibrosis, chronic wounds, and even cell death^25^. Chronic wounds (such as those associated with diabetes) can arise from the persistence of a pro-inflammatory environment driven by continuous ROS release that prevents advancement to the later remodeling stages of wound healing^53,54^. This suggests that misregulation of ROS levels and timing of ROS accumulation following injury can disrupt wound repair outcomes. Indeed, exposure to silver nanoparticles during wound healing and regeneration has been found to inhibit wound-induced ROS generation, leading to the early decline of critical regenerative signals, a decrease in the recruitment of immune cells, and a dampened proliferative response^55^. Therefore, this will be an important consideration in the clinical setting, where dressings for acute and chronic wounds can contain silver nanoparticles due to their antibacterial property^56^.

These data add to the growing body of evidence indicating ROS are highly conserved wound-induced signaling molecules humans^22^, suggesting that more investigation into specific wound-related ROS signaling pathways is needed. We found that distinct signaling programs were associated with differential ROS thresholds. Lower levels of ROS induced only *jun-1* expression as part of wound healing, but were not sufficient to initiate signaling needed for regeneration (Fig. 8B and E). However, higher levels induced expression of wound healing-associated *jun-1*, as well as the expression of *hsp70* required for tissue regeneration (Fig. 8D-E). Importantly, these data revealed that even within a narrow range, different relative levels of ROS can alter gene expression programs, producing radically different morphological outcomes that could have drastic consequences for survival (Fig. 10B). As such, we propose that investigation of potential dose-dependent effects will be vital going forward, simultaneously adding to our understanding of the complex roles of ROS in tissue repair and regeneration.

While we show that ROS are not necessary for the wound site muscle contractions that occur immediately following injury, we found that ROS do regulate wound closure via control of cytoskeletal movements. These findings are consistent with reports that ROS can oxidize actin-binding proteins, as well as actin itself, to promote actin polymerization, cell migration, and cell spreading^57^. Additionally, ROS can also initiate actin cytoskeletal changes through regulation of MAPK signaling. ROS are able to oxidize and inactivate thioredoxin, an inhibitor of the MAP Kinase Kinase Kinase (MAP3K) apoptosis signal-regulating kinase 1 (ASK1)^58^. If not inhibited, ASK1 promotes the activation of the MAPKs c-Jun N-terminal kinase (JNK) and p38, whose transcriptional targets both include the stress-induced transcription factor c-Jun^59^. Decades of research have linked MAP3Ks, JNK, and c-Jun with regulation of the cytoskeleton and actin polymerization (F-actin), and there are many routes by which each can promote cell elongation and migration^60-62^. This pathway has also been linked to wound healing, and in particular the overexpression of c-Jun in diabetic rats was found to accelerate the rate of wound closure^63^.

The literature links between ROS, c-Jun, and wound healing prompted us to investigate this potential relationship in planarians, where our data reveal that ROS are required for wound site expression of *jun-1*. We show that *jun-1* inhibition impairs actin-mediated epithelial stretching during wound healing, phenocopying the impaired healing observed following ROS inhibition. In *Drosophila*, dorsal closure requires the c-Jun homolog (*D-jun*) for the maintenance (but not the initiation) of cell elongation, without which cells readopt a polygonal cell shape, resulting in failed dorsal closure^64^. Additionally, c-Jun knockout mice are born with open eyes and defects in epidermal wound healing, both due to a lack of proper epithelial cell elongation and migration^65^. Thus, regulation of cell elongation via changes in actin polymerization appears to be a highly conserved role of c-Jun.

Together, our data indicate a mechanistic link between injury-induced ROS signaling and wound site *jun-1* expression, which together promote the cytoskeletal changes required for wound closure. However, since we show that ROS regulated wound edge reorganization while *jun-1* did not, it seems clear that ROS signaling during wound healing must include other downstream effectors. This emphasizes the need for further research into this area. For instance, identification of *jun-1*’s DNA binding sites may shed light on the mechanisms that promote epithelial cell motility during planarian wound healing. Activator protein-1 (AP-1), a transcriptional dimer comprised of c-Jun and c-Fos, directly activates genes such as epidermal growth factor receptor (EGFR)^66^. Upstream of EGFR, AP-1 has been shown to activate the cytoskeletal regulators Rac and Rho^61^. EGFR is known to be activated early during planarian regeneration, and its loss has been shown to affect ROS accumulation^40^. Interestingly, EGF signaling is itself known to be upstream of AP-1 activation^67^, suggesting the possibility of a regulatory feedforward mechanism.

Another area for future study will be in the specific type of ROS that endogenously regulate wound repair mechanisms. Our investigations used a general ROS inhibitor (DPI); however, several ROS (such as H_2_O_2_ and superoxide) have been implicated in wound healing and regeneration^52,68^. Additionally, while our data show that application of exogenous H_2_O_2_ was sufficient to rescue wound healing, it is unclear if this occurs through specific redox mechanisms. While ultimately outside the scope of this study, this still leaves many questions unanswered. Do specific ROS independently mediate different downstream signaling programs resulting in wound healing versus regeneration? Does one single ROS, or perhaps the same combination of ROS, mediate both outcomes simultaneously through different mechanisms? Is the origin of ROS after injury extracellular (such as originating from damaged cells at the wound site) or internal through established redox and/or mitochondrial mechanisms? Answers to these questions will be required for more precise manipulation of the appropriate ROS (and downstream signaling) during therapeutic development.

The regulation of ROS levels has implications for therapies beyond those of tissue repair. ROS signaling plays a role in the immune system response, for example where neutrophils release superoxide to control bacterial infections^69^. Careful fine-tuning of ROS levels is also required in stem cells, where differences in concentration can directly impact cell fate decisions such as self renewal, stemness, differentiation, and senescence^70^. ROS is equally critical during aging. For instance, imbalances in ROS levels can cause oxidative damage to chondrocytes, resulting in degradation of cartilage and development of osteoarthritis^71^. Other common and debilitating age-related diseases, such as atherosclerosis, type-2 diabetes, neurodegenerative diseases (like Alzheimer’s), and cancers, have all been linked to redox imbalances, highlighting how furthering our understanding of ROS signaling could have diverse applications for a variety of clinically relevant diseases^72^. Furthermore, it may be beneficial to develop post-injury ROS exposure guidelines, given that H_2_O_2_ is often used to cleanse wounds at concentrations that have been shown to inhibit regeneration^8^. However, the diverse roles of ROS can confound therapeutic approaches, where the presence of numerous redox-sensitive signaling targets and antioxidant control mechanisms can result in unsuccessful attempts to manipulate ROS levels to specific effect. Our results indicate that much more information about ROS signaling during tissue repair is required. Understanding discrepancies between ROS signaling pathways will be critical to the development of targeted treatments for many disease states and improved clinical outcomes.

## Materials and Methods

### Animal care and amputations/injuries

An asexual clonal line of *Schmidtea mediterranea* (CIW4) was maintained at 18°C in the dark. All planarians were kept in worm water; worm water consists of Instant Ocean salts (0.5 g/liter) in ultrapure water of Type 1. Animals were fed no more than once a week with “natural” (no antibiotics or hormones) liver paste made from whole calf liver (Creekstone Farms, Arkansas, KS). Liver was frozen and thawed only once before feeding animals. Worms were starved at least 1 week before experimentation. Animal sizes were as follows: wound healing assays and actin/nuclei labeling, 5-7 mm; *in situ* hybridization and blastema morphology, 3-4 mm; *jun-1* RNAi, 7-8 mm (final size). *S. mediterranea* were amputated via scalpel cuts done under a dissecting microscope on a custom-made cooling Peltier plate, as previously described^73^. For the wound healing assays and ROS accumulation timeline, animals were bisected and resulting head fragments were kept to observe posterior-facing wounds. For actin and nuclei labeled animals, anterior dorsal wounds were made using a hypodermic needle (Air-Tite 27GX2) pushed fully through the worm three times. For *jun-1* gene expression in Fig. 6: experiments used bisected animals as described above. In Fig. 8: healing wounds were performed as to not induce blastema formation as previously described^74^ and regenerating wounds were bisected, except for RNA interference (Fig. 8E), which were amputated above and below the pharynx and observed at the anterior blastema.

### ROS indicator dye assay

The cell-permeant fluorescent general oxidative stress indicator dye, 5-(and-6)-chloromethyl-2′,7′ dicholorodihydrofluorescein diacetate (CM-H2DCFDA; Molecular Probes C6827), was used to visualize ROS accumulation (excitation, 470 nm; emission, 525 nm). One hour before imaging, worms were incubated in 25 μM CM-H2DCFDA made from 10 mM dimethyl sulfoxide (DMSO) stock. For the timeline, fragments that required ROS detection under 1 hour were placed in dye intact, amputated, placed back in dye and imaged at their respective times. After the 1 hour CM-H2DCFDA incubation period, all worms were rinsed three times in fresh worm water and the ventral side was imaged using 35 mm FluoroDishes (WPI FD35-100) and 25 mm round no. 1.5 coverslips (WPI 503508). Signal intensity at the wound site was normalized to signal intensity of the central body to control for differences in dye loading between animals. For Fig. 6A, signal intensity at the wound site is normalized to body brightness. For Fig. 1B: 15 min n=16; 30 min n=16; 45 min n=15; 60 min n=15. For Fig. 1E1: 2 repeats of n≥5, total n=14. For Fig. 1E2: head fragments had 3 repeats of n=5, total n=15; tail fragments had 3 repeats of n≥2, total n=12. For Fig. 1E3: head fragments had 2 repeats of n≥4, total n=9; tail fragments had 2 repeats of n=5, total n=10. For Fig. 1E4: head fragments had 2 repeats of n=5, total n=10; tail fragments had 3 repeats of n≥4, total n=14. For Fig. 1F: Fig. 1F1 had 2 repeats of n=6, total n=12; Fig. 1F2 had 2 repeats of n=5, total n=10. For Fig. 9A: Fig. 9A1 had 2 repeats of n≥5, total n=11; Fig. 9A2 total n=11; Fig. 9A3 had 2 repeats of n≥7, total n=15; Fig. 9A4 had 2 repeats of n≥4, total n=10. For Fig. 9B: healing wounds were from Fig. 9, A1-A2, total n=22; regenerating wounds were from Fig. 1, E2 and E3 and Fig. 9, A3-A4, total n=66.

### Wound healing assay and pharmacological treatments

Wound healing assays were performed as previously described^41^, where open wounds are characterized by the emergence of parenchymal tissue at the wound site. For Fig. 1A: 15 min n=44; 30 min n=44; 45 min n=44; 60 min n=110. ROS accumulation was inhibited with diphenyleneiodonium chloride (DPI; Sigma D2926). For the dose response, animals were presoaked in the given DPI concentration (made from a 3 mM DMSO stock) for 24 hours before amputation and drug was refreshed after amputation. For Fig. 2C: DMSO controls had 11 repeats of n≥5, total n=108; 1 μM DPI had 4 repeats of n≥5, total n=34; 5 μM DPI had 4 repeats of n≥5, total n=34; 10 μM DPI had 5 repeats of n≥5, total n=49; 15 μM DPI had 6 repeats of n≥5, total n=53; 20 μM DPI 4 reps, n≥5, total n=34; 25 μM DPI had 5 repeats of n≥5, total n=50; 35 μM DPI had 6 repeats of n≥5, total n=52; 50 μM DPI had 5 repeats of n≥5, total n=43. All other ROS inhibition experiments used 15 μM DPI to expose intact animals 24 hours in advance and refreshed drug after wounding until scoring; except for H_2_O_2_ rescue experiments where DPI was washed out at injury, rinsed 3 times in worm water, and then placed in the given concentration of H_2_O_2_ (Sigma 7722-84-1) or worm water for controls until scoring. Due to the confounding variable of toxicity, animals that died were excluded from analyses. For Fig. 2D: DMSO controls at 15 min, n=37; 30 min n=37, 45 min n=37, 60 min n=108. For Fig. 2E: DPI at 15 min n=33, 30 min n=33, 45 min n=33, 60 min n=67. For H_2_O_2_ dose response (Fig. 3B): controls (worm water) had 11 repeats of n=10, total n=110; while all H_2_O_2_ concentrations had 4 repeats of n≥10 with 50 μM total n=52, 100 μM total n=53, 200 μM total n=52, 300 μM total n=55, 400 μM total n=54, 500 μM total n=57, 750 μM total n=60, 1000 μM total n=61. For Fig. 3D: controls n=59, control plus H_2_O_2_ n=30, DPI n=49, DPI plus H_2_O_2_ n=30. For Fig. 9F: Controls had 8 repeats of n≥7, total n=69; DPI had 8 repeats of n≥7, total n=62; DPI + 200 μM H_2_O_2_ had 4 repeats of n≥10, total n=45; DPI + 300 μM H_2_O_2_ had 6 repeats of n≥7, total n=56; DPI + 400 μM H_2_O_2_ had 2 repeats of n≥15, total n=37.

### RNA interference

RNAi was performed via feeding of in vitro–synthesized double-stranded RNAi, as previously described^75^. A 518 base pair region of *S. mediterranea jun-1* (SMU15011552) was used to generate *jun-1* RNAi; primers: 5′-AAAGATGAAGTTGTAAAGCA and 3′-AACGACGCTAACTTTAAGTGG. For actin and nuclei labeling experiments: worms were fed with RNAi 4 times over 14 days before being amputated on day 15. For blastema morphology analysis: worms were fed with RNAi 3 times over 9 days before being amputated on day 10. Control RNAi was double-stranded RNA to Venus-GFP, which is not present in the planarian genome. For Fig. 8E: Control RNAi had 2 repeats of n≥10, total n=25; *jun-1* RNAi total n=10; *hsp70* RNAi total n=15. *hsp70* RNAi was made as previously described from the same region as the riboprobe^38^.

### *Whole-mount actin and nuclei labeling/* In situ *hybridization*

Whole mount actin (Phalloidin-FITC; Sigma P5282) labeling was based on a protocol previously described^76^ by killing/demucousing animals in 7.5% NAC (Sigma A7250) in 10 mL 1X PBS for 7.5 min, then fixing with 4% formaldehyde solution (Supelco, FX0410) in 1X PBS for 20 min. After washing 10 min in 1X PBS, samples were permeabilized/blocked in 1X PBS with 0.3% Triton X and 5% horse serum for 1-2 hours. Samples were washed for 15 min in 1X PBS, then incubated overnight at 4°C in 0.5 μM FITC-phalloidin diluted in 1X PBS. The following day samples were washed 3 times for 10 min with 1X PBS, then counterstained for nuclei (Hoechst 33342 10 μg/mL; ThermoFisher H3570) for 10 min. Samples were washed 3 times for 10 min in 1X PBS, then cleared in 90% glycerol/water solution for several hours before mounting and viewing under a confocal microscope. Fluorescence *in situ* hybridization was carried out as previously described^77^. Riboprobe for *jun-1* was generated from the same region as the RNAi. A 552 base pair region of *S. mediterranea hsp70* (SMU15039086) was used to generate riboprobe; the region was from 5′-GGTTTTTGATTTGGGTGGTG to 3′-AGCTGTTGCTATGGGAGC.

### Image collection

All images except actin/nuclei labeling were taken using a ZEISS V20 fluorescence stereomicroscope with AxioCam MRc or MRm camera and ZEN lite software (ZEISS). Fragments were imaged while fully extended and moving to ensure the absence of any tissue bunching, which could affect analyses. Heat maps for visualizing intensity of ROS levels were generated using the standard rainbow lookup table within the ZEN lite software. For actin/nuclei-labeled samples, images were taken on a Nikon C2+ scanning laser confocal microscope. Z-stack imaging was used for all actin/nuclei photos except epithelial stretching, which was observed on a single plane. Adobe Photoshop was used to orient, scale, and improve clarity of images (but not for fluorescent images). All fluorescent images from a single assay (the controls and treated) were taken at the same exposure and magnification settings. Data were neither added nor subtracted; original images are available upon request.

### Quantification and statistical analyses

Significance was determined with a two-tailed Student’s *t* test with unequal variance (using Microsoft Excel) for all, except for the following exceptions: Data from the wound healing assays were analyzed by a two-sample *t*-test between percents (two-tailed) using the Statistics Calculator software (StatPac, V. 4.0). Significance for the ROS timeline (Fig. 1D) and H_2_O_2_-rescued blastema growth (Fig. 9G) was determined using a one-way analysis of variance (ANOVA) with Tukey’s multiple comparison test (using GraphPad Prism version 9.00 for Mac). For Fig. 1D: ANOVA: F-value=15; degrees of freedom (DF) between columns=3; total DF=61. For Fig. 9G: ANOVA: F-value=20.945; DF between columns=4; total DF=95. For ROS indicator dye assays, the magnetic lasso tool was used to obtain mean gray intensity values at the wound site, as well as baseline mean signal intensity values (from the center of the same animal); signal intensity was expressed as wound site mean signal intensity – baseline mean signal intensity, to control for differences in dye loading. For *in situ* hybridization, the magnetic lasso tool was used to obtain mean gray intensity values at the wound site and center of animals; values were expressed as a ratio of wound site mean signal intensity/center mean signal intensity. For Fig. 8B: wound site gene expression was measured on the injured site and compared to the opposing intact region on the same animal. For blastema size, the magnetic lasso tool in Adobe Photoshop was used to generate total pixel counts for both anterior and posterior blastemas (as well as the entire regenerate); to control for worms of different sizes, blastema sizes were expressed as a ratio of blastema size/total regenerate size. For Fig. 9G: Controls had 3 repeats of n≥6, total n=28; DPI had 3 repeats of n≥6, total n=19; DPI + 200 μM H_2_O_2_ had 3 repeats of n≥3, total n=19; DPI + 300 μM H_2_O_2_ had 3 repeats of n≥2, total n=18; DPI + 400 μM H_2_O_2_ had 2 repeats of n≥4, total n=12.

## Acknowledgements

This work was supported by a National Science Foundation grant (#1652312) to W.S.B and an American Association of University Women Fellowship to A.V.VH.

## Author contributions

This work was supervised by W.S.B., who supplied materials and reagents. A.V.VH. conceived of the initial idea and wrote the initial manuscript. A.V.VH. analyzed all the data, with contributions from W.S.B. A.V.VH. performed all the assays, with the exception of: J.M.G. assisted with the wound healing and oxidative stress dye assays, and H_2_O_2_ rescue studies; S.J.H. assisted with RNAi and *in situ* hybridization; L.J.K. assisted with the regeneration rescue imaging and oxidative stress dye timeline. A.V.VH. and W.S.B. prepared the figures. W.S.B. and S.J.H. edited the draft; W.S.B., A.V.VH., S.J.H., and L.J.K. provided input on the final version.

## Competing interests

All authors declare no competing interests.

## Data availability

All relevant data supporting the key findings of this study are available within the article or from the corresponding author upon reasonable request.

## Notes

### Competing Interest Statement

The authors have declared no competing interest.

